# A molecular barcode and online tool to identify and map imported infection with *Plasmodium vivax*

**DOI:** 10.1101/776781

**Authors:** Hidayat Trimarsanto, Roberto Amato, Richard D Pearson, Edwin Sutanto, Rintis Noviyanti, Leily Trianty, Jutta Marfurt, Zuleima Pava, Diego F Echeverry, Tatiana M Lopera-Mesa, Lidia Madeline Montenegro, Alberto Tobón-Castaño, Matthew J Grigg, Bridget Barber, Timothy William, Nicholas M Anstey, Sisay Getachew, Beyene Petros, Abraham Aseffa, Ashenafi Assefa, Awab Ghulam Rahim, Nguyen Hoang Chau, Tran Tinh Hien, Mohammad Shafiul Alam, Wasif A Khan, Benedikt Ley, Kamala Thriemer, Sonam Wangchuck, Yaghoob Hamedi, Ishag Adam, Yaobao Liu, Qi Gao, Kanlaya Sriprawat, Marcelo U Ferreira, Alyssa Barry, Ivo Mueller, Eleanor Drury, Sonia Goncalves, Victoria Simpson, Olivo Miotto, Alistair Miles, Nicholas J White, Francois Nosten, Dominic P Kwiatkowski, Ric N Price, Sarah Auburn

## Abstract

Imported cases present a considerable challenge to the elimination of malaria. Traditionally, patient travel history has been used to identify imported cases, but the long-latency liver stages confound this approach in *Plasmodium vivax*. Molecular tools to identify and map imported cases offer a more robust approach, that can be combined with drug resistance and other surveillance markers in high-throughput, population-based genotyping frameworks. Using a machine learning approach incorporating hierarchical FST (HFST) and decision tree (DT) analysis applied to 831 *P. vivax* genomes from 20 countries, we identified a 28-Single Nucleotide Polymorphism (SNP) barcode with high capacity to predict the country of origin. The Matthews correlation coefficient (MCC), which provides a measure of the quality of the classifications, ranging from −1 (total disagreement) to 1 (perfect prediction), exceeded 0.9 in 15 countries in cross-validation evaluations. When combined with an existing 37-SNP *P. vivax* barcode, the 65-SNP panel exhibits MCC scores exceeding 0.9 in 17 countries with up to 30% missing data. As a secondary objective, several genes were identified with moderate MCC scores (median MCC range from 0.54-0.68), amenable as markers for rapid testing using low-throughput genotyping approaches. A likelihood-based classifier framework was established, that supports analysis of missing data and polyclonal infections. To facilitate investigator-lead analyses, the likelihood framework is provided as a web-based, open-access platform (vivaxGEN-geo) to support the analysis and interpretation of data produced either at the 28-SNP core or full 65-SNP barcode. These tools can be used by malaria control programs to identify the main reservoirs of infection so that resources can be focused to where they are needed most.

## Background

The last three World Malaria Reports have revealed a disturbing rise in malaria cases, and, outside Subsaharan Africa, an increasing proportion of malaria due to *Plasmodium vivax*, undermining the painstaking efforts to reduce transmission over the past decade^1^. These trends highlight the urgent need for new surveillance tools, with greater attention to the non-falciparum species. In today’s global climate, human populations are highly mobile, with imported cases of malaria confounding local control efforts and enhancing the risks of drug resistance spread and outbreaks. There is thus a critical need to develop tools that can help to determine where patients acquired their infection.

The challenge of imported infections is particularly pertinent for *P. vivax*, in view of the parasite’s ability to form dormant liver stages (hypnozoites) that can reactivate weeks to months after the initial infection, as well as highly persistent, low density blood-stage infections ^2,3^. The re-emergence of *P. vivax* in multiple regions where it was once almost eliminated serves as an important reminder of the need to maintain diligent surveillance of this species^4^. In low endemic settings where the incidence of local infections is declining, the relative proportion of imported cases generally rises, emphasizing the importance for surveillance tools that can identify imported *P. vivax* cases. Traditionally, imported cases have been identified and mapped using information on patient travel history, but the persistent blood stage infections and long-latency liver stages constrain the accuracy of this approach in *P. vivax* infections. Molecular tools to identify and map imported *P. vivax* cases offer an attractive complement to traditional epidemiological tools.

Amplicon-based sequencing has become a favored approach for targeted genotyping of malaria parasites. Using highly parallel sequencing platforms, such as the latest generation of Illumina sequencers, amplicon-based sequencing can be applied at moderate to high-throughput, with high accuracy and sensitivity. These platforms are flexible, allowing iterative enhancement of the Single Nucleotide Polymorphism (SNP) barcodes, which can provide an affordable genotyping approach, amenable to population-based molecular surveillance.

Previous studies have used mitochondrial and apicoplast markers to resolve imported from local *P. vivax* isolates, but the resolution of these organelles is constrained^5–7^. In 2015, a panel of 42 SNPs was identified to facilitate parasite finger-printing and geographic assignment^8^. The proposed 42-SNP Broad barcode was derived from genomic data available from 13 isolates from 7 countries. In the last 4 years, the repository of genomic data on *P. vivax* has expanded greatly, allowing further refinement of a parsimonious and widely applicable genotyping barcode.

The primary objective of our study was to develop molecular tools for identifying and characterizing imported *P. vivax* cases amenable to population-based surveillance frameworks, so that these data can be used to inform strategic decisions on where and how to deploy malaria control interventions. We tailored our molecular tools primarily to surveillance frameworks using Illumina or other high-throughput genotyping platforms. As a secondary objective, we sought to identify single gene regions permissible to lower throughput approaches for use in settings or situations where high-throughput or centralized approaches are not feasible. In addition, we provide informatics tools to support users in analyzing genotyping data produced at the barcode that can accommodate missing data and polyclonal infections.

## Materials and Methods

### Overview of the marker selection approach

A flow diagram outlining the steps involved in the marker selection process is provided in Figure 1a. In accordance with the multiplexing features of the Illumina platform, we sought to identify approximately 50 new SNP-based markers to append to the recently published Broad barcode^8^, to provide a composite panel with ≤100 markers for country-level geographic assignment of *P. vivax* infections. The decision to append markers to the Broad barcode rather than selecting a de novo panel of SNPs was pragmatic, aimed at promoting consensus and continuity with existing molecular tools available to the vivax community. A likelihood-based classifier approach was chosen for the respective evaluation of marker sets and end-user data analysis tasks. This approach was chosen since it allows manual addition of specific SNP sets, such as the Broad barcode. Two selector algorithms, hierarchical FST (HFST) and decision-tree (DT), were implemented in the likelihood-based classifier framework to select SNPs with high country-level prediction values. The primary SNP selection method was the HFST selector, which leverages on the prior knowledge of the population structure to inform on a relatively parsimonious SNP set with moderately high prediction. The DT selector, the secondary method, was used to select additional SNPs to append to the Broad barcode and the HFST panel for further enhancement of geographical prediction. A 10-fold cross-validation strategy was used to assess the performance of the selectors with the likelihood-based classification framework.

**Figure 1.**
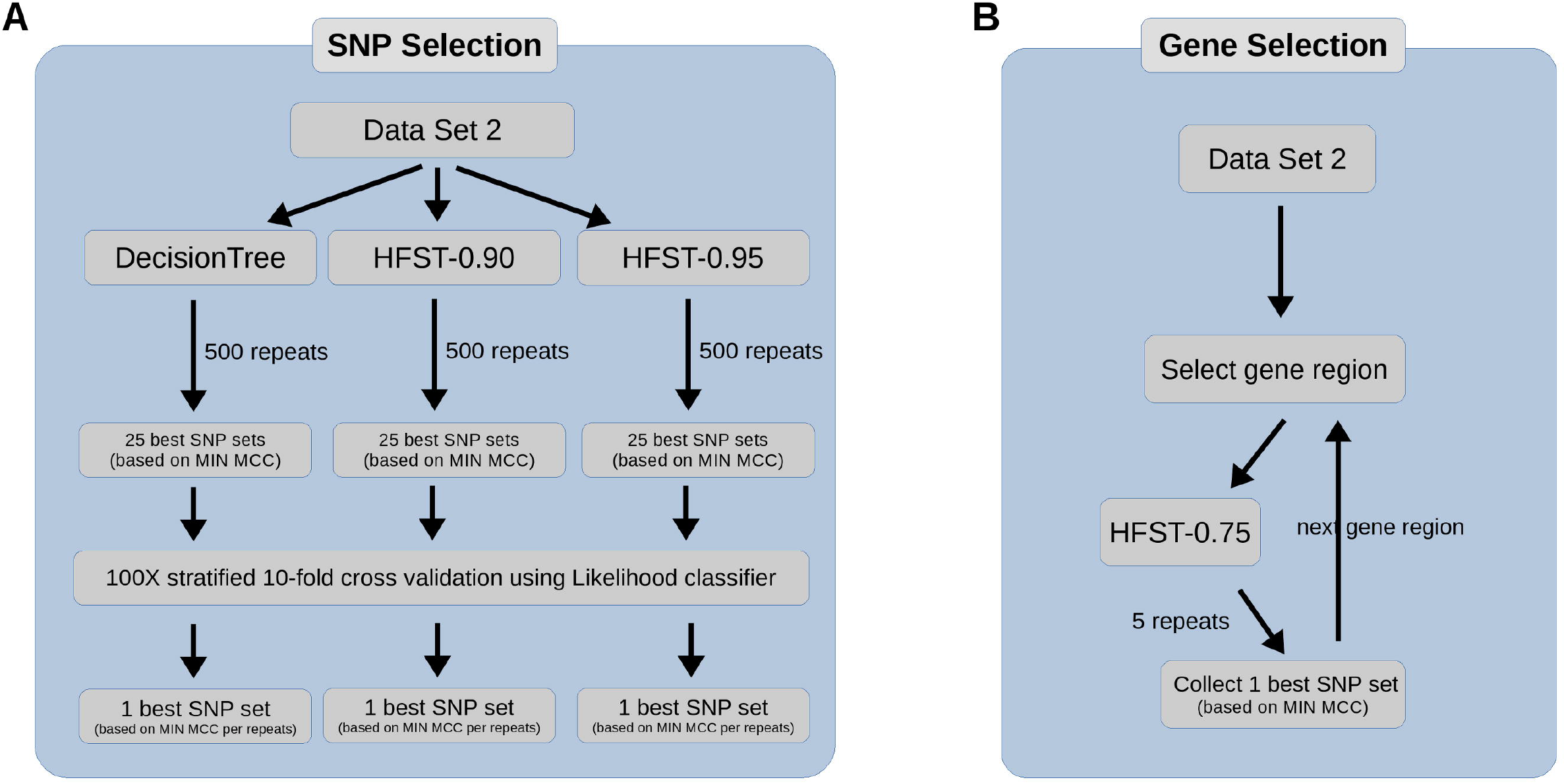
Overview of the marker selection approaches. Flow diagrams illustrating the datasets, selector models and classification approaches used to identify and evaluate independent SNP panels (A) and single gene regions (B). Decision Tree (DT), HFST-0.90 (HFST and DT with FST threshold of 0.90), HFST-0.95 (HFST and DT with FST threshold of 0.95) and HFST-0.75 (HFST and DT with FST threshold of 0.75) represent the SNP selector models. The DT, HFST-0.9 and HFST-0.95 SNP selector models were run in 500 repeats for SNP selection (A), and the HFST-0.75 model was run in 5 repeats for gene selection (B). For SNP selection (A), the top-25 SNP sets were selected from each model for a further 100 repeats of stratified cross-validation from which one SNP set was selected from each of the DT, HFST-0.9 and HFST-0.95 SNP selector models.

To achieve the secondary objective of the study, identifying single gene regions with moderate-to-high country-level resolution, simulations were run across individual genes using the HFST-0.75 (HFST with FST threshold of 0.75) selector model with the likelihood classifier. The top 20 genes with the highest pooled median Matthew Correlation Coefficient (MCC) scores for each selector model were reported (Figure 1b).

### Overview of the web-based data analysis and sharing platform

To establish accessible informatics tools to support users to analyze and interpret their data, a platform was created incorporating data classification tools for determining the most likely country of origin of a sample using genetic data at a given barcode. Existing source code, developed for a microsatellite-based *P. vivax* data sharing and analysis platform ^9^, was modified to create a new web-based platform (vivaxGEN-geo), to collate SNP data generated at the geographic barcode. A likelihood-based classifier approach was chosen for geographic assignment within the vivaxGEN-geo platform owing to the ability to i) incorporate manual SNP sets, ii) evaluate barcodes with missing data, and iii) evaluate heterozygous genotype calls.

### Likelihood-based classifier framework

The custom classifier was developed to handle bi-allelic heterozygote calls for mix-infection cases by treating the samples as diploid samples, as well as missing data by treating as heterozygote calls. The classifier was derived from Bernoulli Naive Bayes with modification to the likelihood equation and elimination of prior probability, since the distribution of our dataset did not reflect the distribution in nature, but rather the implication from sample and extracted DNA quality, as well as the characteristics of the original study such as duration and type of the study. Hence the classifier only depends on the likelihood of the SNP data. The likelihood equation was modified to handle the bi-allelic data as follows:

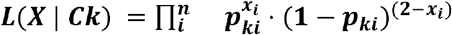

where ***X*** is the SNP data set of a sample, *Ck* is a group (or a country), *x_i_* is the number of alternate alleles at position *i* and *p_ki_* is the frequency of the alternate allele at position *i* of country *k* counted as diploid samples.

### SNP Selection

To select optimal SNPs for country classification, a combination of the HFST and DT selector methods were employed. The DT selector utilized the Python-based scikit-learn package for the decision tree implementation, which employed an optimized version of the CART (Classification And Regression Tree) algorithm and Gini coefficient. To avoid overfitting, a minimum of 3 samples was required for a leaf node. The Hierarchical FST (HFST) approach worked by traversing across a bifurcating guide tree and selecting SNP with the highest FST between the two populations represented by the two nodes of the branch with the assumption that the SNP with the highest FST might differentiate those two populations. If the highest FST of a certain branch was lower than a given threshold during guide tree traversal, the DT method was employed to obtain additional SNPs to separate the given branch. To avoid overfitting, a maximum tree depth of 2 was set for this particular DT step. The HFST analysis in this study was undertaken using a guide tree constructed using Nei’s population distance matrix implemented with a neighbor-joining algorithm.

The classification performance was measured with MCC for each country ^10^. In addition, the pooled median, mean, minimum and first-quartile MCC were collected as additional measurements.

Three models, HFST-0.90 (HFST and DT with FST threshold of 0.90), HFST-0.95 (HFST and DT with FST threshold of 0.95), and pure DT were trained with the full dataset. For each of the three models, 500 repeats were run to allow for different random seeds of the DT analysis, and the top 25 SNP sets with the highest aggregate minimum MCC score as evaluated by the likelihood classifier were obtained from each model. A stratified 100 repeat, 10-fold cross-validation was run on each of the 25 SNP sets from each model, and the best SNP set from each of model, as indicated by highest aggregate minimum MCC score within a repeat, was selected. To compare the Broad SNP panel to the three new SNP panels identified by the HFST-0.90, HFST-0.95 and pure DT selectors, a 500 repeat, stratified 10-fold cross-validation was undertaken on each SNP panel.

### Missing data evaluation

To assess the durability of prediction performance of the SNP sets with missing data, a simulation was run by removing genotype data randomly. The Likelihood classifier was trained against the selected SNP sets using all samples. For each country, 25 samples were sampled randomly with replacement and missing genotype calls were added to the SNP sets in 10%, 20% and 30% proportions. The random samples were then subjected to the trained classifier. This process was run in 100 repeats and MCC-score of the prediction for each country was reported.

### Datasets

The analysis included genomic data on *P. vivax* isolates collected from 26 countries. Published data were included from 19 countries derived from the European Nucleotide Archive^11–15^, and new data from 10 countries (Supplementary Table 1, Supplementary Figure 1). New genomic data were derived from patients recruited to partner studies in Afghanistan, Bangladesh, Bhutan, Colombia, Ethiopia, Indonesia, Iran, Malaysia, Sudan, and Vietnam. With the exception of Colombia, the patient sampling frameworks have been described previously ^11,12,14,16–20^. The samples from Colombia were collected within the framework of cross-sectional epidemiological surveys conducted between 2013 and 2017. Whole genome sequencing, read alignment and variant calling were undertaken within the framework of a *P. vivax* community study in the Malaria Genomic Epidemiology Network (MalariaGEN)^21^. Data was derived from an open dataset of *Plasmodium vivax* genome variation comprising 2,671,112 discovered variants across 1,366 isolates (MalariaGEN manuscript in preparation). The data were initially filtered to derive a set of 670,962 high-quality bi-allelic SNPs with VQSLOD score >0, and minimum read depth and minimum minor allele count of 2. Individual genotype calls were defined as heterozygotes based on an arbitrary threshold of a minor allele ratio > 0.1 and a minimum of 2 reads for each allele; all other genotype calls were defined as homozygous for the major allele. A pair of isolates with distance less than 0.0005 (0.05%) were considered non-independent. Amongst non-independent sample pairs, the isolate with the higher percentage of genotype failures was removed from the dataset; this removal process was iterated until all non-independent isolates had been removed from the dataset. The samples and SNPs were then subjected to further filtering to eliminate missing data using information derived from a simulation which calculated the total number of SNPs with no missing data and the number of consecutive informative SNPs as defined by SNPs with minimum minor allele count (MAC) >2. The remaining dataset was defined as Dataset 1.

The isolates in Dataset 1 were initially assigned to national groups based on the country in which the patient presented at the clinic with the infection. The national-level groupings were evaluated further using country-level assignments derived from the genome-wide data classification with the likelihood classifier. Infections presenting with country classifications differing from the country of presentation were considered suspected imported infections and removed to produce Dataset 2.

Of the 42 Broad barcode SNPs, 37 SNPs were present in the 670K dataset (bi-allelic high-quality SNPs) and exhibited successful amplicon-based sequencing assays (personal communication, Wellcome Sanger Institute Core Sequencing Facility); these 37 SNPs were not present in dataset 1 or 2. A new dataset (Dataset 3) was prepared for evaluation of the Broad barcode comprising of samples with complete data across the 37 Broad barcode SNPs and partial data across SNPs selected from the HFST and DT algorithms.

### Software and Web Service Availability

All custom, in-house scripts used for data filtering, simulation, analyses and visualization are available from http://github.com/trmznt/vivaxgen-geo. The VivaxGEN-geo web service provides a user-friendly online tool for country classification with all SNP sets, and is accessible at https://geo.vivaxgen.org/. The likelihood classifier provided on the online platform has been trained with 809 samples (dataset 4), consisting of all samples with complete data at all SNP sets. The classifier tool reports the three highest likelihoods for country of origin and their associated probabilities.

### Ethics

All samples were collected with written informed consent from patients or their legal guardians. Ethical approvals for the published samples are detailed in the original papers ^11–15^. Approvals for the newly represented studies are outlined in Supplementary Document 1.

## Results

### Geographic clustering patterns using the genome-wide dataset

The primary dataset (Dataset 1) was derived using the missing data simulation, to minimise genotype failures (Supplementary Figure 3), it comprised 854 high-quality samples and 294,628 high-quality informative SNPs, with no missing data. The median percentage of heterozygous calls in each country ranged from 0.02% to 0.08%. Details on the geographic locations of the samples in dataset 1 are presented in Supplementary Table 1. Neighbor-joining analysis on dataset 1 revealed distinct geographic clustering of most countries (Supplementary Figure 4). Exceptions included the isolates from Afghanistan, Iran, India and Sri Lanka, which appeared to form a single cluster, warranting further analysis of this geographic region with larger sample sets. Although several isolates in border regions including Vietnam relative to Cambodia, and Thailand relative to Myanmar, overlapped between countries, the majority of isolates in these countries could be differentiated by national boundaries. Visual inspection of the neighbour-joining tree revealed potential imported cases. Using country-level assignments derived from the genome-wide data classification with the likelihood classifier and manual confirmation of the neighbor-joining tree, 21 isolates presented country classifications differing from the country of presentation (Supplementary Table 1). After exclusion of the imported cases, and countries represented by a single sample, a total of 831 isolates and 20 countries remained, constituting Dataset 2 (Supplementary Table 1).

### Performance of the Broad barcode, HFST and DT SNP selection

When the HFST selector was applied with an FST threshold of 0.90 (HFST-0.90), a set of 28 SNPs (listed in Supplementary Table 2) were identified. This dataset exhibited median MCC scores exceeding 0.9 in all countries with the exception of Vietnam (0.75) and Cambodia (0.80). On increasing the FST threshold to 0.95 (HFST-0.95), the HFST model identified 51 SNPs (listed in Supplementary Table 3), which displayed MCC scores exceeding 0.95 in all countries except for Vietnam (0.85) and Cambodia (0.87). Using the DT selector alone, 50 SNPs (listed in Supplementary Table 4) displayed comparable performance to the 51-SNP panel, with a slightly lower aggregate minimum MCC score.

The results of cross-validation of the classification performance of the five SNP panels (37-SNP Broad barcode, 28-SNP, 28-SNP plus Broad barcode (65-SNP), 50-SNP and 51-SNP panels) are illustrated in Figure 2, and the MCC and F scores reflecting the consensus results of the cross-validation are summarized in Table 1. The performance, ranked from lowest to highest, was: 37-SNP Broad barcode (median MCC = 0.82), 28-SNP (MCC = 0.99), 50-SNP (MCC = 1.00), 65-SNP (MCC = 1.00), and 51-SNP (MCC = 1.00).

**Figure 2.**
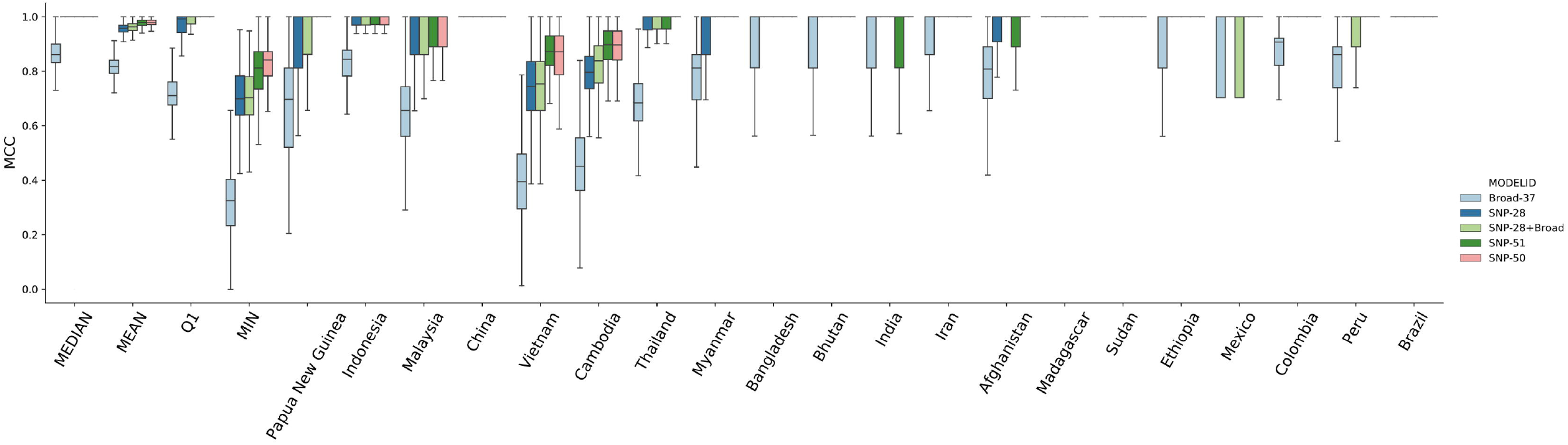
Comparison between the 37-SNP Broad barcode, new marker panels and combined SNP sets. The Broad-37 SNP set reflects 37 of the 42 Broad SNPs represented amongst the 294K high-quality SNPs. The SNP-28 SNP set reflects 28 high-performance SNPs derived from the HFST selector with FST threshold of 0.9. The SNP-28+Broad SNP set reflects the combined Broad-37 and SNP-28 SNP sets for a total of 65 SNPs. The SNP-50 set reflects the 50 SNPs selected by the Decision Tree selector. The SNP-51 set reflects 51 high-performance SNPs from the HFST selector with threshold FST of 0.95. The boxplots present the MCC scores from 500 repeats with stratified 10-fold cross validation for each SNP set. Country labels are provided on the y-axis; MEDIAN, MEAN, Q1 (1st percentile) and MIN reflect the respective summary statistics for the pooled MCC scores across all countries.

**Table 1.** Summary of MCC and F-scores from the consensus results of 500 repeats of the stratified 10-fold cross-validation of the SNP panels

| Population | 37-SNP (Broad) |  | 28-SNP |  | 65-SNP |  | 50-SNP |  | 51-SNP |  |
| --- | --- | --- | --- | --- | --- | --- | --- | --- | --- | --- |
|  | MCC | F | MCC | F | MCC | F | MCC | F | MCC | F |
| Afghanistan | 0.852 | 0.857 | 0.987 | 0.988 | 1 | 1 | 0.974 | 0.975 | 0.987 | 0.988 |
| Bangladesh | 0.796 | 0.778 | 0.881 | 0.875 | 1 | 1 | 1 | 1 | 1 | 1 |
| Bhutan | 0.665 | 0.615 | 1 | 1 | 1 | 1 | 0.894 | 0.889 | 1 | 1 |
| Brazil | 0.815 | 0.8 | 1 | 1 | 1 | 1 | 1 | 1 | 1 | 1 |
| Cambodia | 0.456 | 0.518 | 0.78 | 0.809 | 0.813 | 0.837 | 0.893 | 0.907 | 0.898 | 0.911 |
| China | 0.912 | 0.909 | 1 | 1 | 1 | 1 | 1 | 1 | 1 | 1 |
| Colombia | 0.895 | 0.903 | 1 | 1 | 1 | 1 | 1 | 1 | 1 | 1 |
| Ethiopia | 0.929 | 0.931 | 1 | 1 | 1 | 1 | 1 | 1 | 1 | 1 |
| India | 0.714 | 0.706 | 1 | 1 | 1 | 1 | 0.935 | 0.933 | 0.925 | 0.923 |
| Indonesia | 0.819 | 0.857 | 0.971 | 0.978 | 0.984 | 0.988 | 0.987 | 0.99 | 0.99 | 0.993 |
| Iran | 0.865 | 0.857 | 0.925 | 0.923 | 1 | 1 | 1 | 1 | 1 | 1 |
| Madagascar | 0.894 | 0.889 | 1 | 1 | 1 | 1 | 1 | 1 | 1 | 1 |
| Malaysia | 0.614 | 0.627 | 0.923 | 0.927 | 0.949 | 0.95 | 0.962 | 0.963 | 0.962 | 0.963 |
| Mexico | 0.92 | 0.919 | 1 | 1 | 0.92 | 0.919 | 1 | 1 | 1 | 1 |
| Myanmar | 0.529 | 0.48 | 0.835 | 0.824 | 0.881 | 0.875 | 0.935 | 0.933 | 1 | 1 |
| Papua New Guinea | 0.616 | 0.6 | 0.888 | 0.889 | 0.934 | 0.933 | 0.976 | 0.977 | 0.975 | 0.976 |
| Peru | 0.839 | 0.846 | 1 | 1 | 0.961 | 0.962 | 1 | 1 | 1 | 1 |
| Sudan | 1 | 1 | 1 | 1 | 1 | 1 | 1 | 1 | 1 | 1 |
| Thailand | 0.681 | 0.725 | 0.971 | 0.975 | 0.985 | 0.988 | 0.99 | 0.992 | 0.985 | 0.987 |
| Vietnam | 0.389 | 0.446 | 0.714 | 0.74 | 0.733 | 0.757 | 0.861 | 0.872 | 0.866 | 0.876 |
| **Mean** | **0.76** | **0.763** | **0.944** | **0.946** | **0.958** | **0.96** | **0.97** | **0.972** | **0.979** | **0.981** |
| **Median** | **0.817** | **0.823** | **0.994** | **0.994** | **1** | **1** | **0.995** | **0.996** | **1** | **1** |
| **Q1** | **0.653** | **0.624** | **0.914** | **0.915** | **0.945** | **0.946** | **0.955** | **0.956** | **0.983** | **0.984** |
| **Min** | **0.389** | **0.446** | **0.714** | **0.74** | **0.733** | **0.757** | **0.861** | **0.872** | **0.866** | **0.876** |

### Missing data simulations

In the missing data simulations, genotyping failures had the greatest impact on the classification of samples from Cambodia and Vietnam (Figure 3). With 10% missing data, the median MCCs of the 28-SNP panel exceeded 0.9 in all countries, with exception of Vietnam (MCC = 0.80) and Cambodia (MCC = 0.77). With this level of missing data, the addition of the 37 Broad SNPs (65-SNP panel) increased the median MCC to 0.83 in Vietnam and 0.82 in Cambodia. When missing data increased to 30%, the 65-SNP panel achieved median MCCs above 0.9 in most countries, with exception of Vietnam (MCC = 0.79) and Cambodia (MCC = 0.75). The 50- and 51-SNP panels both achieved MCC scores exceeding 0.95 for all countries except Cambodia (0.80-0.82) and Vietnam (0.83-0.85) with 10-30% missing data.

**Figure 3.**
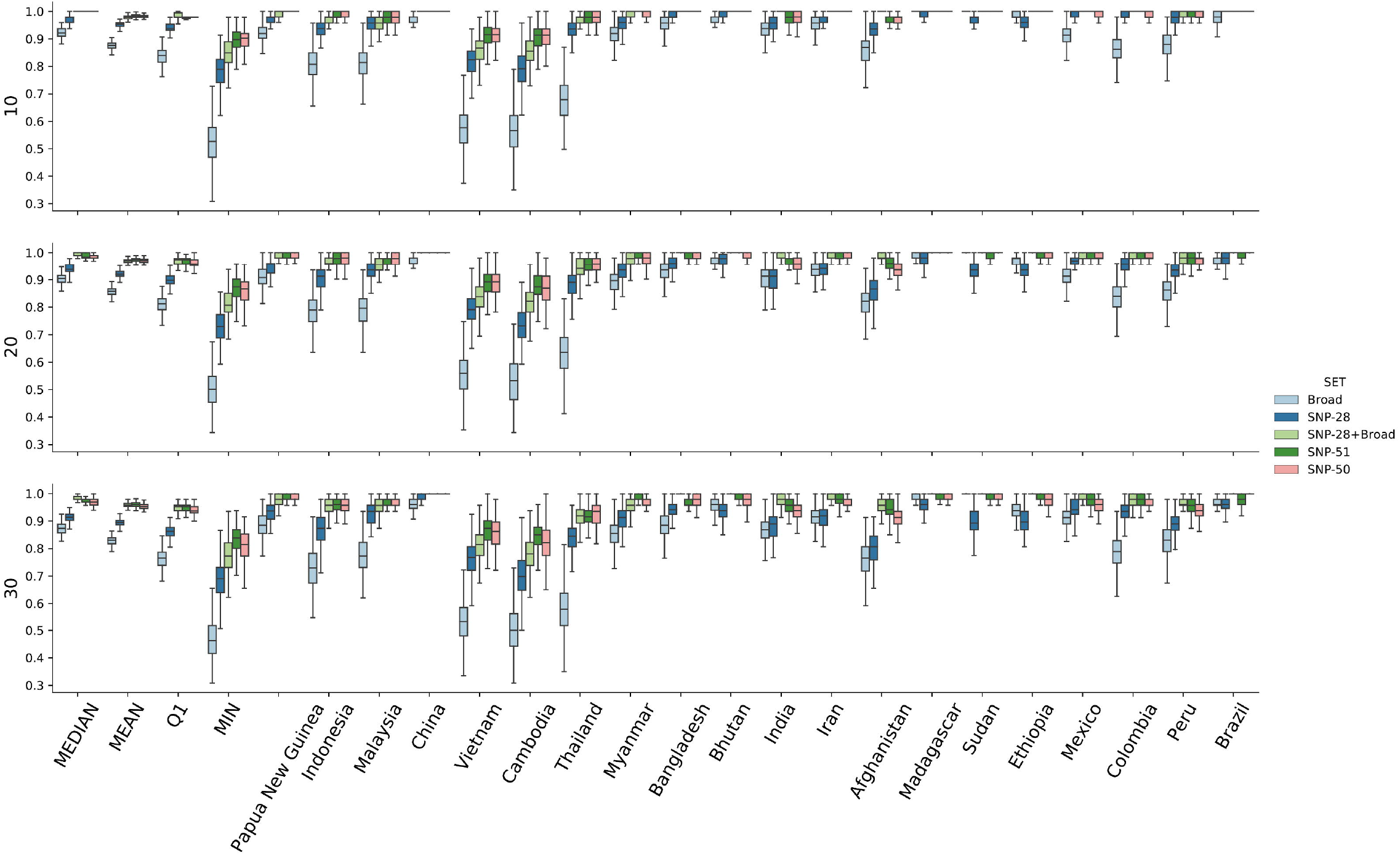
Simulation of missing data in the 28-SNP, 65-SNP, 50-SNP and 51-SNP panels. Result of 200 repeats, 25 samples per country simulation of missing data (genotype fails) of 10%, 20% and 30% against the 37-SNP Broad barcode, 28-SNP set, 65-SNP set (28-SNP + Broad panel), 51-SNP and 50-SNP set. The 65-SNP set demonstrated marginally better performance relative to the 28-SNP set with missing data. However, both the 50-SNP and 51-SNP panels outperformed the 65-SNP panel with missing data.

### Evaluation of single gene regions to predict country classification

The suitability of single genes to predict country classifications were assessed by simulations of individual genes using HFST-0.75 selector model with the likelihood classifier framework. The top 20 genes with the highest pooled median MCC scores for the HFST-0.75 are presented in Supplementary Table 5. The highest prediction capacity, with median MCC score of 0.68, was PVP01_0302600, a gene coding a 11.5 Kb conserved protein with unknown function. The gene list also included three members of the *cysteine repeat modular protein family* (CRMP): CRMP1 (median MCC = 0.63), CRMP3 (MCC = 0.57) and CRMP4 (MCC = 0.56).

## Discussion

The primary objective of the study was to develop molecular tools amenable to population-based surveillance frameworks to identify and map imported *P. vivax* infections. Using machine-learning methods, 3 new SNP panels were identified with high country classification performance, able to distinguish imported *P. vivax* infections across a range of endemic scenarios. The most parsimonious panel, the 28-SNP barcode, exhibited high country classification, and can be appended to the 37 bi-allelic, assayable Broad barcode SNPs for marginal improvement in predictive capacity in samples with moderate levels of missing data. The combined 65-SNP barcode generated robust country classification in most endemic areas, even when the proportion of missing data rose to 30%. However, the validity of the 65-SNP barcode was lower in Cambodia and Vietnam, a likely reflection of the porous border between these two countries. Although the 50- and 51-SNP panels achieved better resolution in these areas, characterization of parasite transmission across borders with high levels of gene flow may be addressed better by the addition of markers suited to an analysis of identity-by-descent^22^. The application and wider validation of the 65-SNP barcode is underway, with amplicon-based sequencing assays already established for the 37 Broad barcode SNPs, and under development for the 28 new markers.

The analysis and interpretation of “real-world” genotyping data raises significant challenges from low-quality samples such as those collected on dried blood spots. In anticipation of these needs we established a likelihood-based framework with the capacity to deal with polyclonal infections as well as missing data. This framework has been integrated into the vivaxGEN-geo online platform, so that users can analyze and interpret their data without needing complex bioinformatics skills and avoiding the subjective visual inspection of neighbour-joining trees or principal component plots. Whilst the informatics tools implemented in vivaxGEN-geo are tailored to *P. vivax*, we anticipate that a similar approach can be adapted to other species. To facilitate wider application the source code will be made publicly available.

The variants in the 28-SNP panel are located in genes representing a range of functions, some of which may be unstable over time. Although our dataset represents one of the most geographically diverse panels of *P. vivax* isolates currently available, with representation of all of the major vivax-endemic regions, the predictive capacity of the derived tools are likely to be constrained by the geographic representation of the reference panel. In particular, representation from central and south America and the Indian subcontinent were limited in our data set. Despite this limitation the dataset used comprises good representation of isolates from areas of public health relevance, including the epicenter of chloroquine-resistant *P. vivax* in Papua Indonesia ^23,24^. The likelihood-based framework is able to re-evaluate the predictive potential of current marker sets as new genomic data become available, so that the selected SNP panels can be refined further in an iterative process. Furthermore, as the reference panel expands with increasing data generated at the barcode SNPs, the accuracy of the likelihood-based classifications will improve.

In addition to the independent selection of SNPs, a number of informative genes were identified, each of which had moderately high geographic resolution power. Genotyping of these genes or gene regions are amenable to standard capillary sequencing, offering an alternative approach, albeit with slightly lower resolution, to define a parasite’s geographic origin in settings where high-throughput genotyping facilities are unavailable. The genes with the greatest geographic resolution, included members of the *cysteine repeat modulator protein* (CRMP) family (CRMP1, CRMP3 and CRMP4) implicated in essential roles in parasite transmission from the mosquito to the human host^25^. It is plausible that the CRMP genes have maintained high geographic differentiation to ensure parasite adaptation to the local vector species. Although adaptations of these genes are likely to temporally stable, the resolution of these loci may be constrained by the distributions of host Anopheles vector species.

In 2017, up to 100% of all confirmed malaria cases in 17 malaria-endemic countries in the Asia-Pacific region, the Middle East and the Americas, where *P. vivax* infections predominate, were reported as being infections ^1^. Malaria control programs in these countries can utilize the information derived from the molecular tools provided from our analysis to assess the efficacy of ongoing interventions in reducing local transmission, and to determine the major routes of infection importation. The tools have potential to reduce ambiguity for certificating malaria elimination by the World Health Organization, where one of the key requirements is the demonstration that all malaria cases detected in-country over at least three consecutive years were imported. For this purpose, countries approaching elimination will need to maintain archival samples for future molecular comparisons against putatively imported cases.

The molecular *P. vivax* geographic classification tools presented are designed to empower users in malaria-endemic countries to analyze and interpret locally produced genotyping data with comparison to globally available datasets. Amplicon-based sequencing of the full 65-SNP barcode is being developed and will be combined with other surveillance markers at central laboratories in endemic partner countries of the Asia Pacific Malaria Elimination Network (www.apmen.org). The data generated from these centers will inform researchers, National Malaria Control Programs and other key stakeholders on the incidence, epidemiology and key reservoirs of imported malaria, and, in doing so, help to target resources to where they are needed most.

## Acknowledgements

We would like to thank the patients who contributed their samples to the study, local health facilities, and the health workers and field teams who assisted with the sample collections. We also thank the staff of the Wellcome Sanger Institute Sample Logistics, Sequencing, and Informatics facilities for their contributions.

---

**Supplementary Figure 1.**
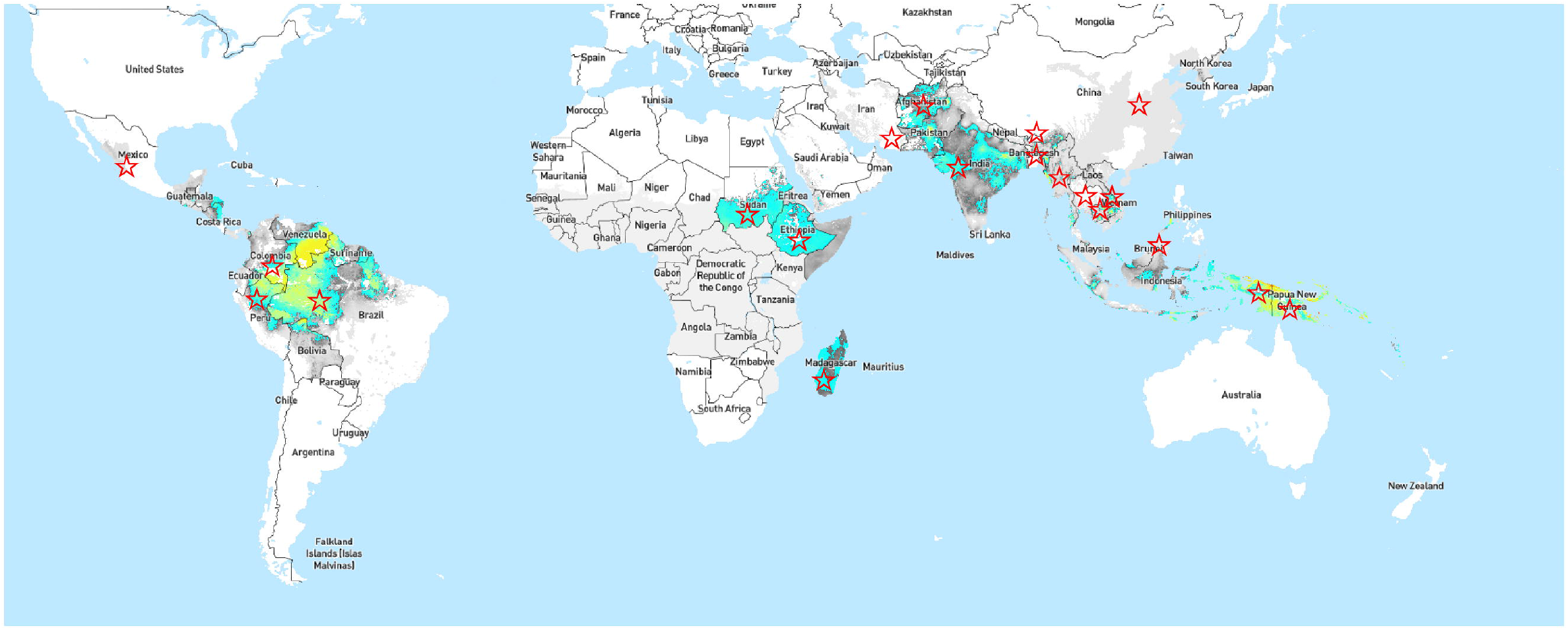
*P. vivax* prevalence map pinpointing the countries included in the study. *P. vivax* prevalence map from the Malaria Atlas Project (*Plasmodium vivax* parasite rate in all ages globally (1-99) from (2000-2017)^26^, with counties included in dataset 2 demarked by stars.

**Supplementary Figure 2.**
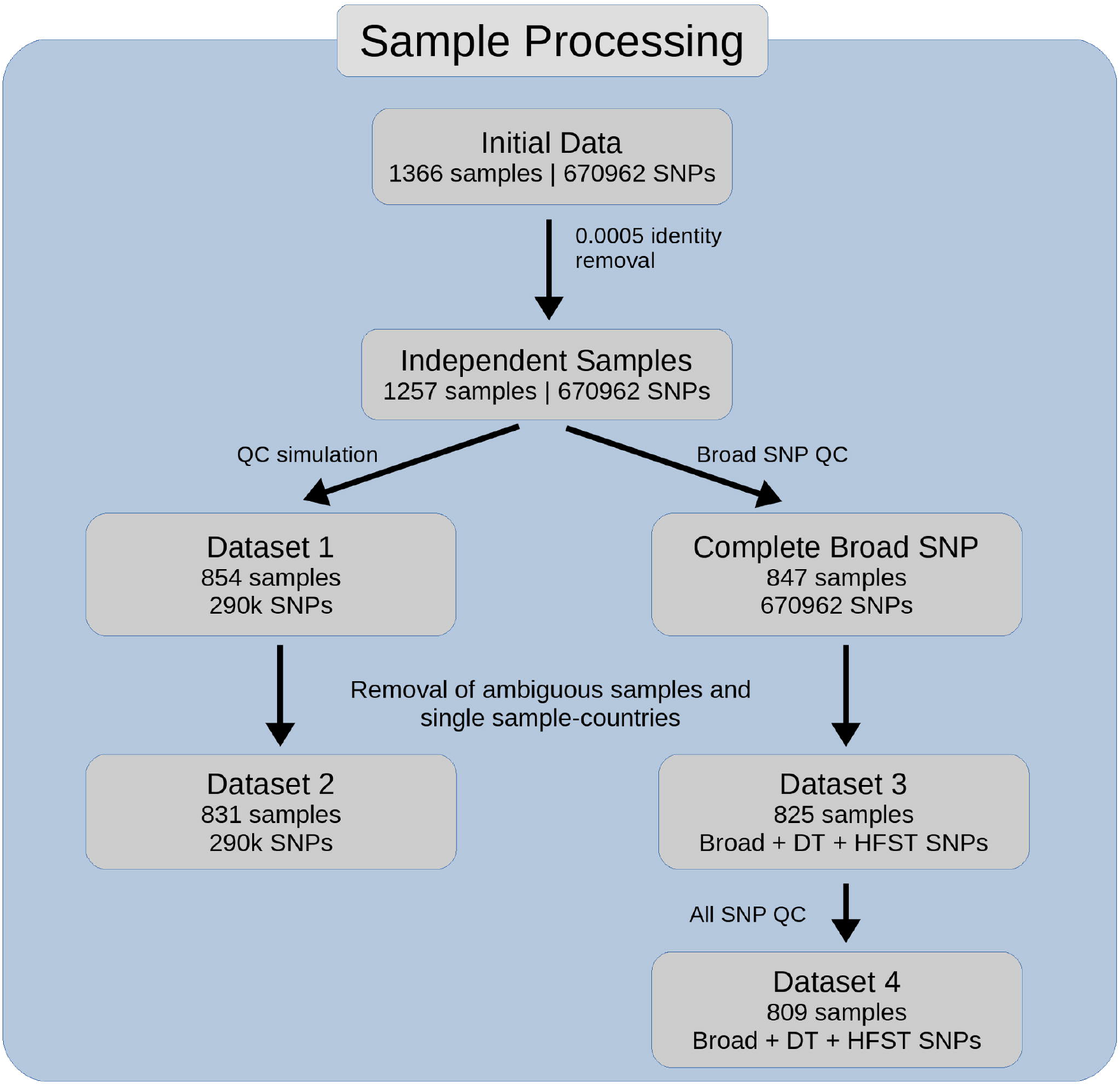
Overview of the datasets

**Supplementary Figure 3.**
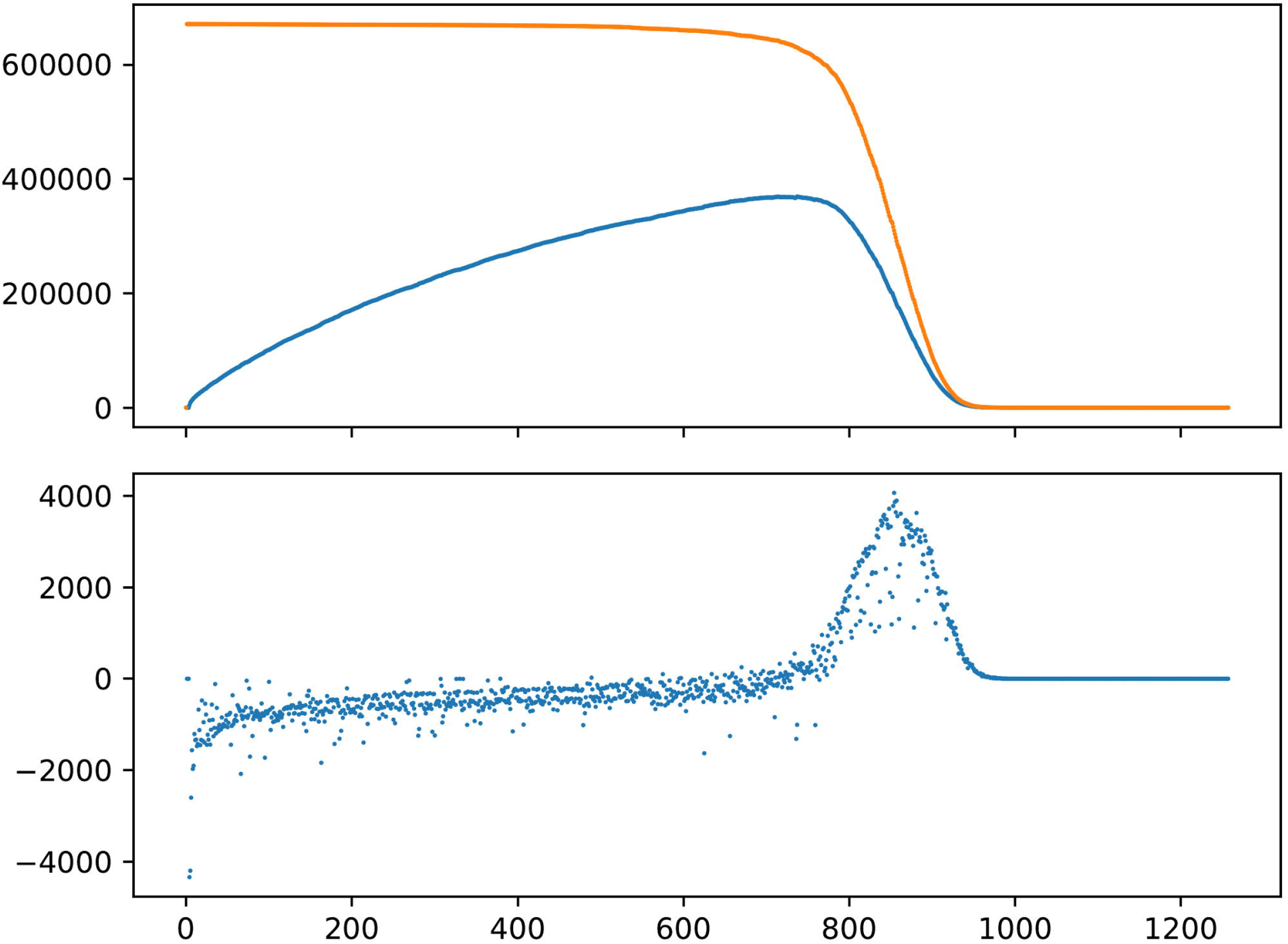
Output from the data quality simulation. The upper panel shows the number of complete SNPs (green), complete informative SNPs with minor allele count (MAC) = 1 (orange) and complete informative SNPs with MAC = 2 (red) against the number of samples. The lower panel shows the number of differences in SNPs between consecutive number of samples, with informative SNPs with MAC = 1 (blue) and informative SNPs with MAC = 2 (orange). The maximum of both MAC=1 and MAC=2 were 958 samples.

**Supplementary Figure 4.**
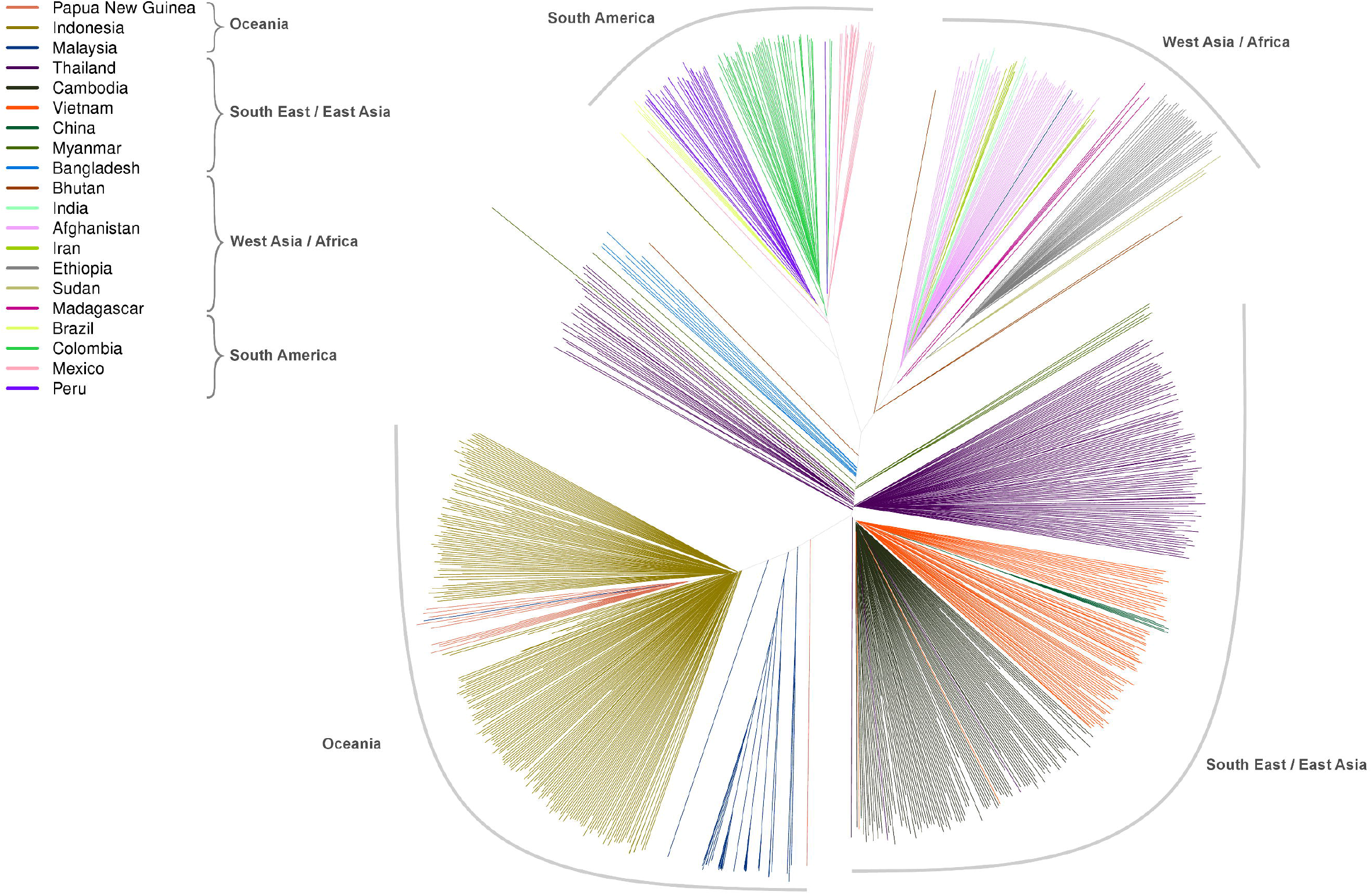
Neighbour-joining tree of the global dataset. The tree was constructed using genotyping data from 854 samples at 294K SNPs.

