## Supplementary Table 1 for "A molecular barcode and online tool to identify and map imported infection with *Plasmodium vivax*"

**Supplementary Table 1: Summary of the patient samples**

| **Population** | **N (Initial)** | **N (independent)** | **N (high quality): dataset 1** | **N (training): dataset 2** | **N (Broad comparison): dataset 3** |
| --- | --- | --- | --- | --- | --- |
| Afghanistan | 250 | 245 | 49 | 49 | 44 |
| Bangladesh | 28 | 28 | 9 | 9 | 8 |
| Bhutan | 9 | 9 | 4 | 4 | 4 |
| Brazil | 25 | 25 | 6 | 6 | 4 |
| Cambodia | 124 | 116 | 111 | 111 | 112 |
| China | 5 | 5 | 5 | 6 | 5 |
| Colombia | 107 | 96 | 58 | 58 | 61 |
| El Salvador | 1 | 1 | 1 | 0 | 0 |
| Ethiopia | 43 | 42 | 33 | 33 | 31 |
| India | 14 | 14 | 7 | 7 | 7 |
| Indonesia | 282 | 252 | 212 | 212 | 214 |
| Iran | 15 | 15 | 9 | 9 | 6 |
| Madagascar | 4 | 4 | 4 | 5 | 4 |
| Malaysia | 109 | 66 | 45 | 45 | 42 |
| Mauritania | 1 | 1 | 1 | 0 | 0 |
| Mexico | 20 | 20 | 20 | 23 | 20 |
| Myanmar | 9 | 9 | 8 | 8 | 8 |
| Nicaragua | 1 | 1 | 1 | 0 | 0 |
| North Korea | 1 | 1 | 1 | 0 | 0 |
| Panama | 1 | 1 | 1 | 0 | 0 |
| Papua New Guinea | 34 | 34 | 24 | 24 | 21 |
| Peru | 47 | 47 | 42 | 42 | 38 |
| Sri Lanka | 1 | 1 | 1 | 0 | 0 |
| Sudan | 13 | 13 | 4 | 4 | 4 |
| Thailand | 129 | 125 | 124 | 124 | 121 |
| Vietnam | 92 | 85 | 74 | 74 | 71 |
| **Total** | **1366** | **1257** | **854** | **831** | **825** |

The initial dataset (N initial) consisted of all patient samples in the MalariaGEN Plasmodium Vivax Community Project release 4 (PV4) with the exception of unpublished data from contributors from non-Menzies partner sites. The independent patient samples (N independent) comprised all infections from the initial dataset (N initial) after filtering out isolates from pairs with nucleotide distance less than 0.0001 (0.01%). The high-quality dataset (N high quality) consisted of the 854 independent patient samples with complete data at the 294,628 high-quality SNPs. The training dataset (N training) consisted of high-quality samples after reassignment based on the neighbor-joining tree. For identification of new markers to supplement the Broad barcode SNPs, the high-quality dataset (N high-quality) was filtered to exclude samples with missing data at the 37 high-quality Broad markers and 28 HFST-derived SNPs.
