## Supplementary Table 2 for "A molecular barcode and online tool to identify and map imported infection with *Plasmodium vivax*"

**Supplementary Table 2: List of the variants in the 28-SNP panel**

| **Chromosome** | **Position** | **Reference allele** | **Alternative allele** | **Gene** | **Amino acid change** |
| --- | --- | --- | --- | --- | --- |
| PvP01_02_v1 | 215587 | C | T | PIP5K | G279R |
| PvP01_04_v1 | 401564 | A | C | ACS9 | K660T |
| PvP01_05_v1 | 160690 | G | T | PVP01_0503800 | M337I |
| PvP01_05_v1 | 576647 | A | G | YIF1 | A775 |
| PvP01_07_v1 | 498997 | G | T | CRMP1 | K1217N |
| PvP01_07_v1 | 510043 | C | T | Intergenic | N/A |
| PvP01_08_v1 | 647036 | T | G | ADA2 | Q1560P |
| PvP01_08_v1 | 1405207 | G | C | PVP01_0832600 | H485Q |
| PvP01_08_v1 | 1506940 | G | A | Intergenic | N/A |
| PvP01_09_v1 | 644970 | G | A | PVP01_0913800 | S5337N |
| PvP01_10_v1 | 189119 | C | A | PPT | G135 |
| PvP01_10_v1 | 480601 | G | A | MDR1 | L845F |
| PvP01_10_v1 | 656127 | G | T | CCR4 | V2369L |
| PvP01_11_v1 | 961323 | C | T | Intergenic | N/A |
| PvP01_11_v1 | 1548740 | G | T | Intergenic | N/A |
| PvP01_12_v1 | 98366 | T | G | Intergenic | N/A |
| PvP01_12_v1 | 2198756 | C | T | PVP01_1253800 | N/A |
| PvP01_12_v1 | 2779908 | T | C | Intergenic | N/A |
| PvP01_12_v1 | 2926892 | C | T | PVP01_1269600 | E569K |
| PvP01_13_v1 | 261941 | C | T | Intergenic | N/A |
| PvP01_13_v1 | 1195172 | C | T | Intergenic | N/A |
| PvP01_13_v1 | 1661666 | A | C | RAP1 | Y234D |
| PvP01_14_v1 | 203651 | A | T | NHE | D917E |
| PvP01_14_v1 | 770449 | C | T | Intergenic | N/A |
| PvP01_14_v1 | 1270401 | G | C | PPPK-DHPS | A553G |
| PvP01_14_v1 | 1270911 | C | G | PPPK-DHPS | G383A |
| PvP01_14_v1 | 1313512 | A | G | JmjC1 | N/A |
| PvP01_14_v1 | 1434209 | C | A | PIGO | F1247L |
