## Supplementary Table 3 for "A molecular barcode and online tool to identify and map imported infection with *Plasmodium vivax*"

**Supplementary Table 3: List of the variants in the 51-SNP panel**

| **Chromosome** | **Position** | **Reference allele** | **Alternative allele** | **Gene** | **Amino acid change** |
| --- | --- | --- | --- | --- | --- |
| PvP01_01_v1 | 273223 | T | C | PPM12 | N776D |
| PvP01_02_v1 | 215587 | C | T | PIP5K | G279R |
| PvP01_03_v1 | 764851 | C | T | PVP01_0318000 | A698T |
| PvP01_03_v1 | 823594 | G | A | Intergenic | N/A |
| PvP01_04_v1 | 853944 | C | G | PVP01_0421000 | H1792Q |
| PvP01_05_v1 | 160690 | G | T | PVP01_0503800 | M337I |
| PvP01_05_v1 | 440699 | G | A | NT2 | V186M |
| PvP01_05_v1 | 576647 | A | G | YIF1 | A775 |
| PvP01_05_v1 | 914054 | A | T | Intergenic | N/A |
| PvP01_06_v1 | 645726 | A | G | CLAMP | F446S |
| PvP01_07_v1 | 498997 | G | T | CRMP1 | K1217N |
| PvP01_07_v1 | 510043 | C | T | Intergenic | N/A |
| PvP01_07_v1 | 720618 | G | C | Intergenic | N/A |
| PvP01_08_v1 | 638172 | G | T | Intergenic | N/A |
| PvP01_08_v1 | 737001 | G | A | RIPR | D794N |
| PvP01_08_v1 | 1011465 | C | G | PVP01_0822700 | V241 |
| PvP01_08_v1 | 1467855 | G | A | PVP01_0834700 | L3119 |
| PvP01_08_v1 | 1506940 | G | A | Intergenic | N/A |
| PvP01_09_v1 | 644970 | G | A | PVP01_0913800 | S5337N |
| PvP01_09_v1 | 1101235 | G | A | PVP01_0924700 | H1141 |
| PvP01_09_v1 | 1884013 | T | G | IMC1b | E141D |
| PvP01_10_v1 | 445674 | C | A | PVP01_1010000 | H172Q |
| PvP01_10_v1 | 451770 | T | G | Intergenic | N/A |
| PvP01_10_v1 | 480601 | G | A | MDR1 | L845F |
| PvP01_11_v1 | 503992 | A | T | PVP01_1112200 | F3054I |
| PvP01_11_v1 | 677441 | T | C | PVP01_1115800 | Y133 |
| PvP01_11_v1 | 1548740 | G | T | Intergenic | N/A |
| PvP01_11_v1 | 1846016 | A | G | PVP01_1143300 | L747 |
| PvP01_12_v1 | 304829 | C | T | Intergenic | N/A |
| PvP01_12_v1 | 1430008 | T | C | Intergenic | N/A |
| PvP01_12_v1 | 1449274 | A | T | Intergenic | N/A |
| PvP01_12_v1 | 1695837 | C | A | PVP01_1241500 | M616I |
| PvP01_12_v1 | 1695950 | C | T | PVP01_1241500 | V579M |
| PvP01_12_v1 | 2007831 | A | T | PVP01_1249300 | S449 |
| PvP01_12_v1 | 2353408 | C | A | Intergenic | N/A |
| PvP01_12_v1 | 2441608 | G | C | MDR2 | V43L |
| PvP01_12_v1 | 2926892 | C | T | PVP01_1269600 | E569K |
| PvP01_13_v1 | 610451 | G | C | PVP01_1313400 | L724 |
| PvP01_13_v1 | 1079771 | G | A | Intergenic | N/A |
| PvP01_13_v1 | 1479779 | G | A | Intergenic | N/A |
| PvP01_13_v1 | 1661666 | A | C | RAP1 | Y234D |
| PvP01_14_v1 | 86024 | G | A | PVP01_1401900 | A77V |
| PvP01_14_v1 | 879619 | G | T | Intergenic | N/A |
| PvP01_14_v1 | 1270401 | G | C | PPPK-DHPS | A553G |
| PvP01_14_v1 | 1270911 | C | G | PPPK-DHPS | G383A |
| PvP01_14_v1 | 1313512 | A | G | JmjC1 | N/A |
| PvP01_14_v1 | 1434209 | C | A | PIGO | F1247L |
| PvP01_14_v1 | 2279901 | G | C | PVP01_1452200 | S915T |
| PvP01_14_v1 | 2694648 | T | A | Intergenic | N/A |
| PvP01_14_v1 | 2882227 | G | C | PVP01_1467500 | V595 |
| PvP01_14_v1 | 2972154 | C | T | PVP01_1469500 | R283Q |
