## Supplementary Table 4 for "A molecular barcode and online tool to identify and map imported infection with *Plasmodium vivax*"

**Supplementary Table 4: List of the variants in the 50-SNP panel**

| **Chromosome** | **Position** | **Reference allele** | **Alternative allele** | **Gene** | **Amino acid change** |
| --- | --- | --- | --- | --- | --- |
| PvP01_03_v1 | 121829 | A | C | PVP01_0302600 | G1220 |
| PvP01_03_v1 | 211805 | A | G | PVP01_0304000 | N1497D |
| PvP01_04_v1 | 401564 | A | C | ACS9 | K660T |
| PvP01_04_v1 | 680799 | C | A | PVP01_0416700 | G415 |
| PvP01_04_v1 | 868268 | T | C | Intergenic | N/A |
| PvP01_05_v1 | 576647 | A | G | YIF1 | A775 |
| PvP01_05_v1 | 722002 | G | C | PVP01_0517200 | T435S |
| PvP01_05_v1 | 866159 | A | G | PVP01_0521300 | Y101H |
| PvP01_05_v1 | 911869 | C | G | PVP01_0522600 | D191H |
| PvP01_05_v1 | 1071362 | C | T | PVP01_0526400 | S275F |
| PvP01_05_v1 | 1355394 | G | A | Intergenic | N/A |
| PvP01_06_v1 | 369313 | C | T | PVP01_0608300 | S4223 |
| PvP01_06_v1 | 419653 | G | C | PVP01_0609200 | L456F |
| PvP01_06_v1 | 910609 | G | A | Intergenic | N/A |
| PvP01_07_v1 | 496776 | C | T | CRMP1 | S477L |
| PvP01_07_v1 | 498997 | G | T | CRMP1 | K1217N |
| PvP01_07_v1 | 1155644 | C | A | PVP01_0727200 | C946F |
| PvP01_08_v1 | 664019 | C | A | Intergenic | N/A |
| PvP01_08_v1 | 1011312 | T | C | PVP01_0822700 | Y190 |
| PvP01_08_v1 | 1011465 | C | G | PVP01_0822700 | V241 |
| PvP01_08_v1 | 1023638 | C | T | PVP01_0822800 | P2591L |
| PvP01_08_v1 | 1506827 | A | G | Intergenic | N/A |
| PvP01_09_v1 | 644970 | G | A | PVP01_0913800 | S5337N |
| PvP01_09_v1 | 1069761 | T | G | Intergenic | N/A |
| PvP01_09_v1 | 1884013 | T | G | IMC1b | E141D |
| PvP01_10_v1 | 155145 | T | C | Intergenic | N/A |
| PvP01_10_v1 | 656127 | G | T | CCR4 | V2369L |
| PvP01_10_v1 | 1264983 | C | A | PVP01_1029200 | P3579 |
| PvP01_11_v1 | 320177 | T | G | Intergenic | N/A |
| PvP01_11_v1 | 909011 | T | C | CEPT | F69L |
| PvP01_11_v1 | 963759 | C | T | ACS | L647 |
| PvP01_11_v1 | 1460240 | A | G | ENR | L292 |
| PvP01_11_v1 | 1548740 | G | T | Intergenic | N/A |
| PvP01_12_v1 | 98366 | T | G | Intergenic | N/A |
| PvP01_12_v1 | 353253 | G | T | ALBA2 | N/A |
| PvP01_12_v1 | 882252 | G | A | PVP01_1222500 | S343 |
| PvP01_13_v1 | 261941 | C | T | Intergenic | N/A |
| PvP01_13_v1 | 971555 | G | A | MIT3 | L228 |
| PvP01_13_v1 | 1262094 | G | A | SPB4 | L537 |
| PvP01_13_v1 | 1621952 | A | G | Intergenic | N/A |
| PvP01_13_v1 | 1711446 | T | C | PVP01_1339300 | K1121R |
| PvP01_14_v1 | 1270401 | G | C | PPPK-DHPS | A553G |
| PvP01_14_v1 | 1270911 | C | G | PPPK-DHPS | G383A |
| PvP01_14_v1 | 1313512 | A | G | JmjC1 | N/A |
| PvP01_14_v1 | 1315135 | A | G | JmjC1 | N/A |
| PvP01_14_v1 | 1434209 | C | A | PIGO | F1247L |
| PvP01_14_v1 | 1786095 | C | G | PVP01_1441200 | N/A |
| PvP01_14_v1 | 1962674 | C | T | K1 | T677 |
| PvP01_14_v1 | 2646079 | C | T | PVP01_1461200 | S587F |
| PvP01_14_v1 | 2971005 | T | C | PVP01_1469500 | N666S |
