## Supplementary Table 5 for "A molecular barcode and online tool to identify and map imported infection with *Plasmodium vivax*"

**Supplementary Table 5. List of the genes with the top 20 MCC scores**

| **MCC** | **F** | **Gene** | **K** |
| --- | --- | --- | --- |
| 0.682712 | 0.679019 | PVP01_0302600 | 57 |
| 0.670033 | 0.659739 | PVP01_1022500 | 39 |
| 0.63265 | 0.627083 | PVP01_1404700 | 46 |
| 0.628669 | 0.618677 | CRMP1 | 23 |
| 0.614111 | 0.607825 | PVP01_1139900 | 43 |
| 0.610236 | 0.607097 | PVP01_1262900 | 35 |
| 0.60204 | 0.585121 | PVP01_1011500 | 42 |
| 0.585187 | 0.573916 | PVP01_1241000 | 33 |
| 0.573111 | 0.549787 | PVP01_0526800 | 31 |
| 0.572246 | 0.570313 | PVP01_1121400 | 40 |
| 0.571136 | 0.566053 | PVP01_1430700 | 40 |
| 0.56788 | 0.54707 | CRMP3 | 29 |
| 0.563723 | 0.546034 | PVP01_0116000 | 44 |
| 0.562524 | 0.555565 | PVP01_1225000 | 31 |
| 0.559193 | 0.554087 | CRMP4 | 43 |
| 0.55983 | 0.557904 | PVP01_0606900 | 37 |
| 0.558556 | 0.543393 | G377 | 29 |
| 0.546512 | 0.535291 | SET1 | 49 |
| 0.542839 | 0.537559 | ApiAP2 | 36 |
| 0.537995 | 0.533571 | EPAC | 36 |
