## Supplementary Document 1 for "A molecular barcode and online tool to identify and map imported infection with *Plasmodium vivax*"

**Supplementary Document 1. Ethics approvals for the new genomic data**

Ethical approvals for the new genomic were provided by the following ethics boards; Islamic Republic of Afghanistan Ministry of Public Health Institutional Review Board, Afghanistan, The Ministry of Health Evaluation Committee on Ethics in Biomedical Research, Vietnam, Oxford Tropical Research Ethics Committee, UK (1014–13), and the Human Research Ethics Committee of the Northern Territory Department of Health and Menzies School of Health Research, Australia (HREC-13-1991) for sampling in Afghanistan and Vietnam; the Ethics Review Committee of the International Centre for Diarrheal Diseases and Research, Bangladesh (PR-14053) and the Human Research Ethics Committee of the Northern Territory Department of Health and Menzies School of Health Research, Australia (HREC-15-2336) for sampling in Bangladesh; the Research Ethics Board of the Ministry of Health in Bhutan (REBH 2012/031), and the Human Research Ethics Committee of the Northern Territory Department of Health and Menzies School of Health Research, Australia (HREC 2012-1871) for sampling in Bhutan; the Bioethics Committee of the Institute for Medical Research, Faculty of Medicine, University of Antioquia, Colombia (BE-IIM), and the Ethics Committee for Human Research at the Centro Internacional de Entrenamiento e Investigaciones Medicas (CIDEIM), Colombia (08-2015) for sampling in Colombia; the Ethics Committee of the Infectious and Tropical Disease Research Center, Hormozgan University of Medical Sciences (HUMS 9014) for sampling in Iran.
